# Primate-specific microRNA-1202 regulates dopaminergic neurogenesis by targeting APC2 and modulates WNT/β-catenin signaling pathway in midbrain organoid

**DOI:** 10.1101/2025.01.19.633822

**Authors:** Xiaohang Long, Dandan Cao, See-Wing Chan, Suyu Hao, Lingxiao Liu, Wai-Yee Chan, Hoi-Hung Cheung

**Affiliations:** School of Biomedical Sciences, Faculty of Medicine, The Chinese University of Hong Kong, Hong Kong S.A.R., China; Centre for Regenerative Medicine and Health, Hong Kong Institute of Science and Innovation, Chinese Academy of Sciences, China; Shenzhen Key Laboratory of Fertility Regulation, Reproductive Medicine Center, The University of Hong Kong – Shenzhen Hospital, Shenzhen, China; Key Laboratory for Regenerative Medicine, Ministry of Education, School of Biomedical Sciences, Faculty of Medicine, The Chinese University of Hong Kong, Hong Kong S.A.R., China

**Keywords:** miR-1202, dopaminergic neuron, midbrain organoid, APC2, WNT signaling, primate-specific miRNA

## Abstract

MicroRNAs (miRNAs) are generally evolutionarily conserved, but a small number of them are found exclusively in primates. MiR-1202 is a primate-specific miRNA that was previously revealed as being involved in major depression disorder (MDD). Moreover, the genomic locus where miR-1202 locates (6q25.3) is strongly associated with recurrent early-onset MDD and neurodevelopmental disorder. We hypothesize that miR-1202 plays a unique role in the brain in fine-tuning the transcriptional network that governs neurogenesis. Here, we reported that microdeletion of miR-1202 resulted in retarded organoid growth but enhanced dopaminergic (DA) neuron differentiation in midbrain organoids. Integrated analysis of miRNA-interacting targets and transcriptional changes in miR-1202 knockout cells revealed that APC2, a negative regulator of canonical WNT/β-catenin pathway, was a downstream target of miR-1202. Knockdown of APC2 resulted in attenuated DA neurogenesis, in contrast to the enhanced DA neurogenesis in miR-1202 knockout cells. Treatments of the brain organoids with antidepressants increased miR-1202 expression and simultaneously inhibited APC2 expression. Consistently, APC2 expression was found to increase in MDD patients. Our results suggest that miR-1202 regulates dopamine neurogenesis by inhibiting the APC2/WNT signaling pathway, which plays a distinct role in the midbrain-hindbrain regional patterning during the early development of the central nervous system.

## INTRODUCTION

In animals, a small number of genes and non-coding RNAs (ncRNAs) have evolved specifically in primates. The uniqueness of these genes and regulatory RNAs in primates, including humans, suggests a complex developmental program that enhances communication and cognitive functions (An et al., 2023; Florio et al., 2018). Many of these primate-specific genes and ncRNAs remain to be characterized, particularly regarding their relevance to human brain development and diseases (Prodromidou and Matsas, 2019; Xing et al., 2024).

One notable example is hsa-mir-1202 (miR-1202), a primate-specific microRNA (miRNA) that is enriched in the brain. MiR-1202 has been linked to the pathophysiology of major depressive disorder (MDD) (Lopez et al., 2014). In patients with MDD, expression levels of miR-1202 are decreased, while those receiving antidepressant treatment show increased expression in the remission group (Fiori et al., 2017). Additionally, miR-1202 has been identified as a sensitive serum biomarker in multiple studies (Fiori et al., 2017; Gattas et al., 2022; Gheysarzadeh et al., 2018). Interestingly, microdeletions at 6q25.2-25.3, the location of miR-1202, have been observed in patients with mental retardation, microcephaly, and developmental delays featuring dysmorphic characteristics (Lukusa et al., 2001; Nagamani et al., 2009). However, the function of this miRNA in the brain, particularly during human brain development, remains poorly understood.

In this study, we investigated the role of miR-1202 in midbrain development using human pluripotent stem cell (hPSC)-derived brain organoids as a model. Our results revealed that the knockout (KO) of miR-1202 led to enhanced dopaminergic (DA) neurogenesis. Integrated analyses of RNA sequencing (RNA-seq) and RNA pulldown-seq indicated that APC2, a negative regulator of the WNT/β-catenin pathway, is a downstream target of miR-1202. The knockdown of *APC2* resulted in impaired DA neurogenesis. Additionally, treatment of the brain organoids with antidepressants increased miR-1202 expression while simultaneously decreasing *APC2* expression. Notably, *APC2* expression was found to be elevated in patients with MDD. Overall, our results suggest that miR-1202 regulates DA neurogenesis by inhibiting the APC2/WNT pathway, which plays a distinct role in regional patterning and midbrain development during early central nervous system formation.

## RESULTS

### Generation of midbrain organoid for studying dopaminergic differentiation

To investigate the role of miR-1202 in neurogenesis, we generated a microdeletion at the miR-1202 locus in hPSCs. Using CRISPR-Cas9 mediated gene editing, we obtained several KO clones with the entire mature miRNA sequence and a short upstream sequence deleted (**Supplementary Figure 1**). From these, we selected two single-cell-derived KO clones (KO2 and KO3) and a mixed polyclonal KO (KO1) for our experiments.

To mimic neurodevelopment in the brain, we employed a three-dimensional differentiation protocol that allowed the generation of self-organized midbrain-like organoids to study the function of miR-1202 (Jo et al., 2016). In normal wild-type (WT) cells, embryoid bodies formed a neuroepithelial structure by day 9 (D9), following dual-SMAD inhibition using Dorsomorphin and A83-01 in combination with WNT signaling activation by CHIR99021. The addition of SHH agonist and FGF8 induced mesencephalic (midbrain) fate. Afterward, matrigel-embedded organoids were allowed to grow, self-organize, and mature in organoid maturation media (**Figure 1a**). Quantitative RT-PCR (qPCR) analysis confirmed the induction of DA neuron differentiation, as evidenced by the increased expression of markers such as *FOXA2*, *LMX1A*, and *OTX2* from D9 to D44. Additionally, neural transcription factors like *ASCL1* and *NEUROG2*, along with pan-neuronal cytoskeletal proteins such as *TUBB3* (TUJ1) and *MAP2*, were highly expressed starting from D30. The presence of dopamine phenotype was supported by the expression of DA neuron markers including *DDC* (DOPA decarboxylase), *TH* (tyrosine hydroxylase), and *NR4A2* (nuclear receptor subfamily 4 group A member 2) (**Figure 1b**). All these markers demonstrated significant upregulation during midbrain organoid differentiation.

**Figure 1.**
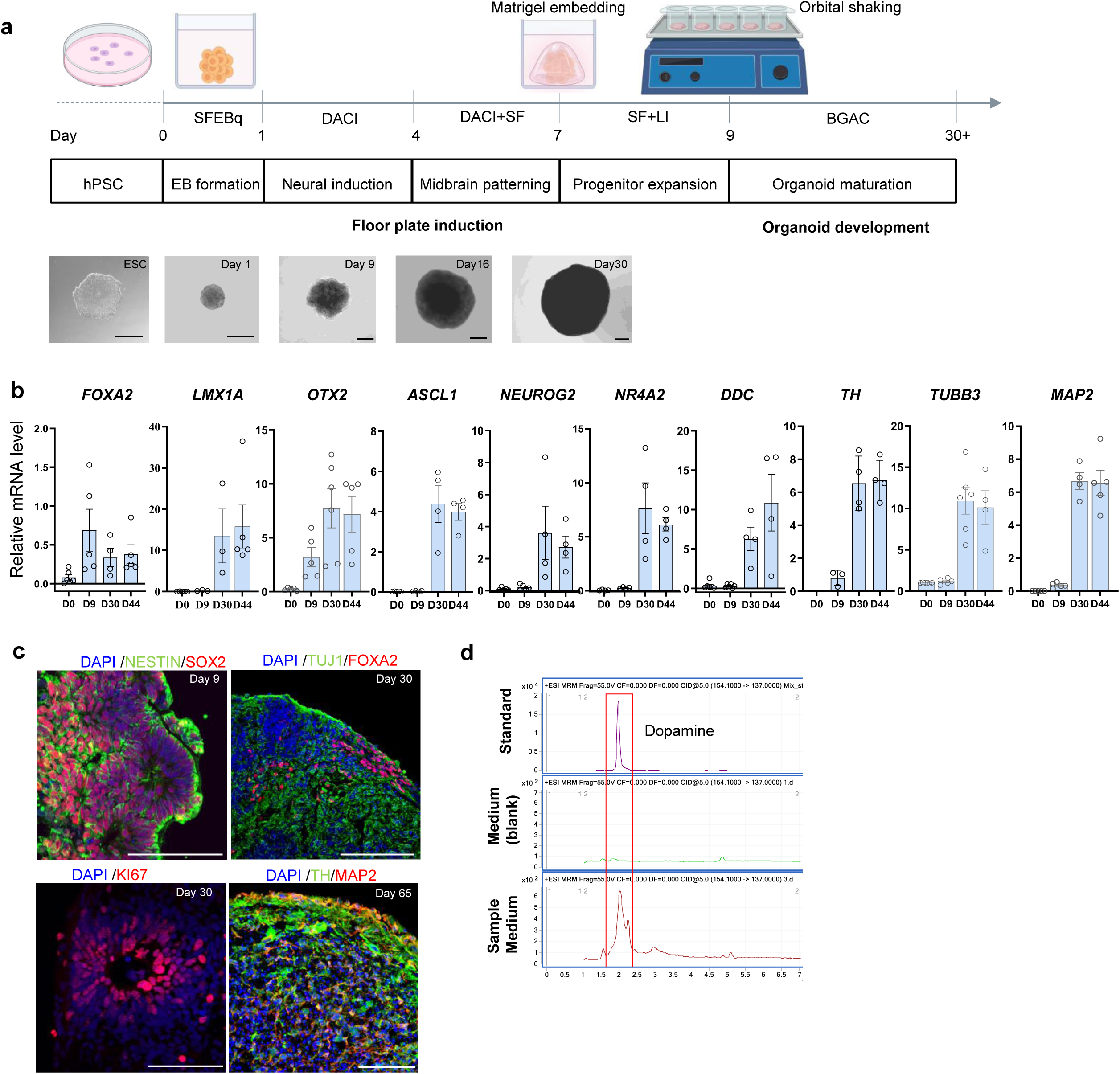
A midbrain 3D organoid model for studying DA neurogenesis. a) Schematic diagram of generating midbrain organoid for studying neuron differentiation. Representative images of 3D organoids were shown (not in scale). SFEBq: serum-free floating culture of embryoid body-like aggregates with quick aggregation; DACI: dorsomorphin, A83-01, CHIR99021 and IWP2; SF: SAG and FGF8; LI: laminin and insulin; BGAC: BDNF, GDNF, ascorbic acid and db-cAMP. Scale bars, 500 µm. b) Q-PCR analysis of DA neuron markers and pan-neuron markers in the organoids during differentiation. c) Immunofluorescent staining of different neural markers in the organoids during differentiation. Scale bars, 100 µm. d) Detection of dopamine in the culture medium by liquid chromatography-mass spectrometry (LC-MS) at D48.

We also performed immunofluorescent staining to verify the expression of representative markers at different stages of the developing organoids: SOX2 and NESTIN at D9, TUJ1 and FOXA2 at D30, and TH and MAP2 at D65. Ki-67 positive cells were observed at D30, indicating that the organoids contained both proliferative progenitor cells and mature neurons (**Figure 1c**). Furthermore, we analyzed dopamine biosynthesis using liquid chromatography-mass spectrometry (LC-MS). The culture medium collected at D48 showed a major peak that overlapped with the dopamine standard, suggesting that the organoids at this stage were capable of synthesizing and releasing the neurotransmitter dopamine (**Figure 1d**).

### Knockout of miR-1202 alters organoid growth and development

Next, we generated midbrain organoids using the miR-1202 KO hPSCs (KO1, KO2, and KO3), along with the WT control cells. At D9, the KO cells formed cell aggregates and neuroepithelial structures similar to those observed in the WT controls (**Figure 2a**). The size and weight of the organoids were comparable between the WT and KO groups at D9 and D16. However, the miR-1202 KO organoids began to show smaller size and lower weight, with these differences becoming more pronounced at D30 and D44 (*p* < 0.001) (**Figure 2b, c**). To investigate whether the smaller size and weight of the KO organoids were due to decreased cell proliferation, we treated the D30 organoids with bromodeoxyuridine (BrdU) to label actively proliferating cells. Flow cytometry analysis indicated a lower percentage of BrdU^+^ cells in all the KO organoids (KO1: *p* < 0.01; KO2: *p* < 0.5; KO3: *p* < 0.5) compared to the WT control (**Figure 2d**). Interestingly, the expression of *MAP2*, a marker associated with more differentiated neurons, was higher in the KO organoids, suggesting a more mature phenotype (**Figure 2e**). Additionally, we performed immunofluorescent staining to assess the expression of the floor plate marker FOXA2 and the midbrain marker OTX2. Surprisingly, despite being smaller in size, the KO organoids exhibited higher levels of FOXA2^+^ and OTX2^+^ cells (**Figure 2f, g**). The upregulation of *FOXA2* and *OTX2* transcripts was further confirmed by qPCR (**Figure 2e**). Since these transcription factors specify progenitor cells toward DA neuron fate, the deletion of miR-1202 appears to promote DA neurogenesis.

**Figure 2.**
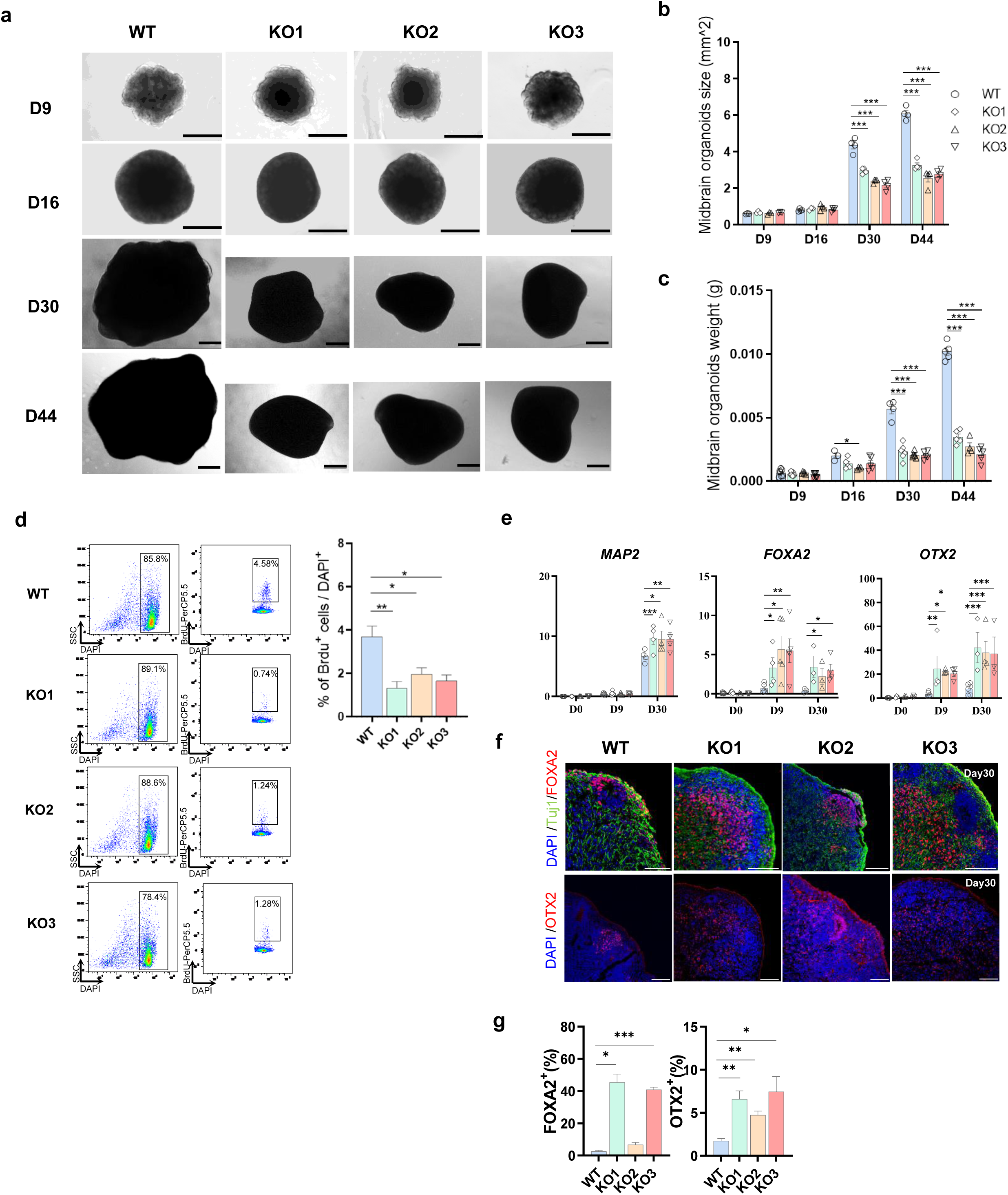
MiR-1202 KO suppresses organoid growth and enhances midbrain transcription factors. a) Representative images of the developing organoids in WT and miR-1202 KO cells (KO1 for mixed clones, KO2 & KO3 for single cell-derived clones). b) Quantification of the organoid size at different time points. c) Quantification of the organoid weight at different time points. d) Flow cytometry analysis of BrdU incorporation for the actively proliferative cells at D30. Quantification was shown. e) Q-PCR analysis of midbrain DA markers (*FOXA2* and *OTX2*) and pan-neuronal marker (*MAP2*). f) Immunofluorescent staining FOXA2, OTX2 and TUJ1 in the D30 organoids. g) Quantification of FOXA2^+^ and OTX2^+^ cells in the D30 organoids. Error bars represent mean ± SEM (b, c, d, e and g).

### Knockout of miR-1202 upregulates genes that promote DA neurogenesis

To further investigate the roles of miR-1202 in the differentiation of DA neurons, we performed RNA-sequencing (RNA-seq) on the WT and miR-1202 KO organoids at two stages: the early progenitor stage at D9 and the more differentiated stage at D30. At D9, we identified 454 upregulated genes and 875 downregulated genes (with a fold change of ≥ 2 or ≤ −2, and adjusted *p*-value < 0.05) (**Figure 3a**). At D30, 349 genes were upregulated and 761 genes were downregulated under the same criteria (**Figure 3b**). Pathway enrichment analysis using Wikipathway and Gene Ontology (GO) Biological Process at both D9 and D30 indicated that the upregulated pathways in KO organoids were primarily associated with DA neurogenesis and neuron differentiation (**Figures 3c,d**). Among the genes that were enriched in the DA neurogenesis pathway and upregulated in KO organoids were transcription factors such as *NKX6-1*, *EN1*, *OTX2*, *FOXA2*, *NKX2-2*, *NEUROD1*, and *NEUROG2*, as well as the Sonic hedgehog gene *SHH*, which promotes floor plate patterning (**Figures 3e,f**). We confirmed the differential expression of specific markers (*LMX1A*, *EN1*, *DDC*, *TH*, and *SHH*) in the KO organoids using qPCR (**Figure 3h**). Additionally, we performed immunofluorescence and Western blot analyses to quantitatively assess the protein levels of TH, the rate-limiting enzyme in dopamine biosynthesis. Both analyses demonstrated consistent increase of TH protein in the KO groups (**Figures 3i,j**). As TH catalyzes the initial step of dopamine synthesis, we measured the dopamine levels in the culture media. Consistently, we observed higher dopamine levels in the KO groups at D48 (**Figure 3k**). Collectively, these transcriptional readout and biomarkers suggest that the depletion of miR-1202 favors DA neurogenesis.

**Figure 3.**
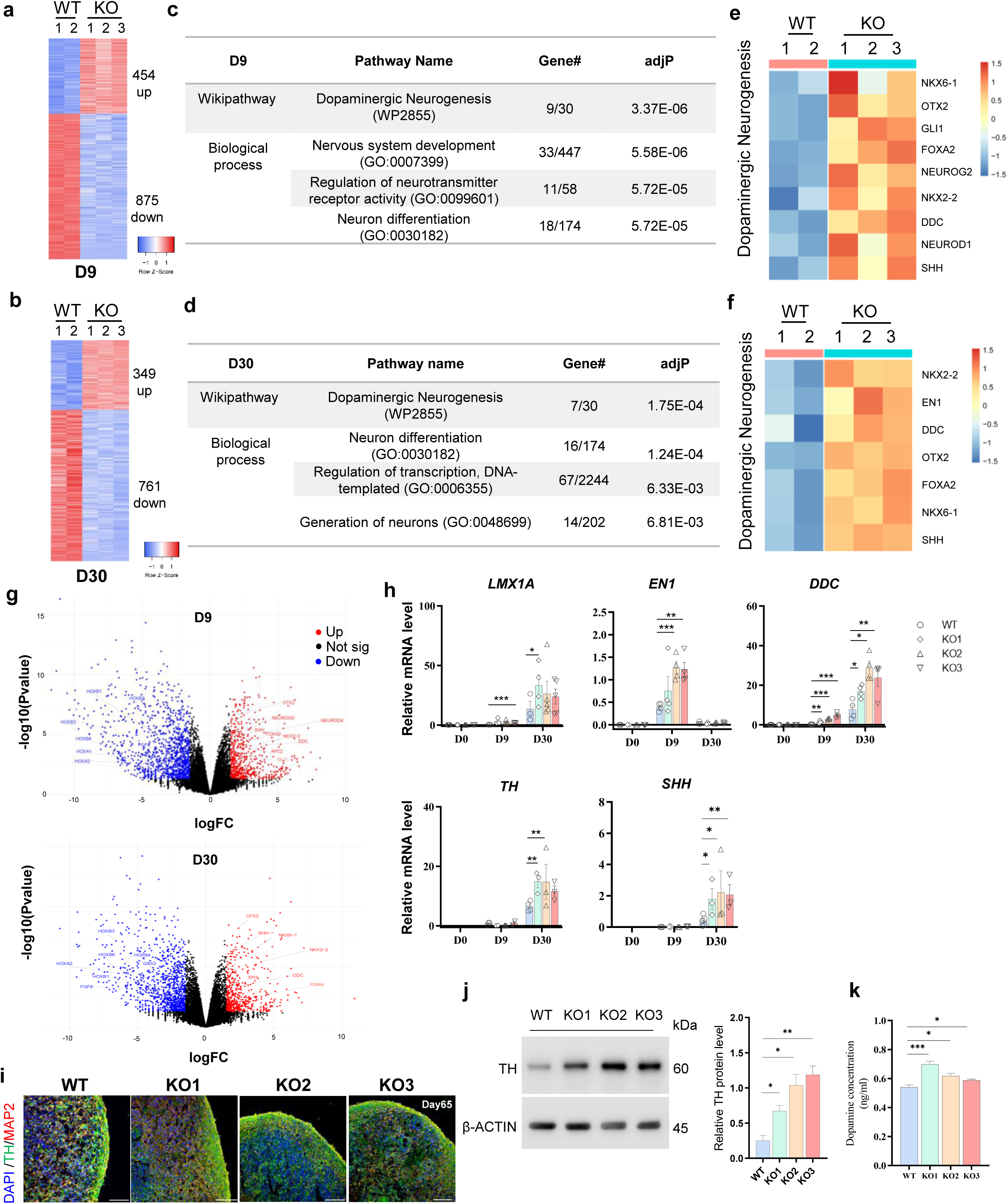
MiR-1202 KO promotes DA neurogenesis and dopamine synthesis. a, b) Heatmaps of the differentially expressed genes (DEGs) in the miR-1202 KO organoids at D9 (a) and D30 (b), respectively. c, d) Pathways enriched for the upregulated DEGs in the miR-1202 KO organoids at D9 (c) and D30 (d), respectively. Gene#, the number of annotated genes in the input list / number of annotated genes in the reference list. *adjP*, *p*-value adjusted by multiple test adjustment. e, f) Lists of the DEGs in the dopaminergic neurogenesis pathway at D9 (e) and D30 (f), respectively. g) Volcano plots showing the significantly upregulated (red) and downregulated (blue) genes at D9 and D30 (fold change > 2, *adjP* < 0.01). h) Q-PCR analysis of the midbrain progenitor transcription factors (*LMX1A* and *EN1*), dopamine synthesis enzymes (*DDC* and *TH*) and signalling molecule (*SHH*). i) Immunofluorescent staining of TH and MAP2 in D65 organoids. Scale bars, 100 µm. j) Western blot of TH protein level in D44 organoids. Quantification of protein amount was shown. k) Dopamine levels in WT and miR-1202 KO organoids were measured by LC-MS. Error bars represent mean ± SEM (h, j and k).

### Knockout of miR-1202 downregulates genes promote hindbrain development

The KO of miR-1202 also resulted in the downregulation of a substantial number of genes: 875 genes at D9 and 761 genes at D30. To understand the implications of this downregulation, we examined the GO terms associated with these genes. We found that there was enrichment in biological processes related to anterior/posterior pattern specification at both D9 and D30 (**Figure 4a**). Among the genes significantly downregulated in the KO cells, we identified homeobox genes expressed in the hindbrain and spinal cord along the anteroposterior axis during brain regionalization, including *HOXA4*, *HOXB4*, and *HOXB6*, which were confirmed by qPCR (**Figure 4b**). Besides homeobox genes, FGF8, the morphogen protein expressed on the hindbrain side of the isthmic organizer, overlaps with GBX2, which opposes OTX2 and promotes hindbrain development (Harada et al., 2016). We thus examined the expressions of *FGF8* and *GBX2*, and found significant downregulations of *FGF8* at D9 and *GBX2* at D9 and D30 (**Figure 4c**). Furthermore, as the Raphe nuclei in the brainstem typically express serotonin receptors that are the targets of antidepressants, we investigated whether the miR-1202 KO affected the expression of these receptors. Notably, the serotonin receptor genes *HTR1A*, *HTR1B*, and *HTR2A* were all significantly downregulated in KO cells (**Figure 4d**). However, the gene expressions of serotonin biosynthesis enzymes *TPH1* and *TPH2*, as well as the transporter *SLC6A4*, did not show consistent changes in the KO groups (**Supplementary Figure 2**). Together, these transcriptional analyses suggest a suppression of the hindbrain development in the KO groups.

**Figure 4.**
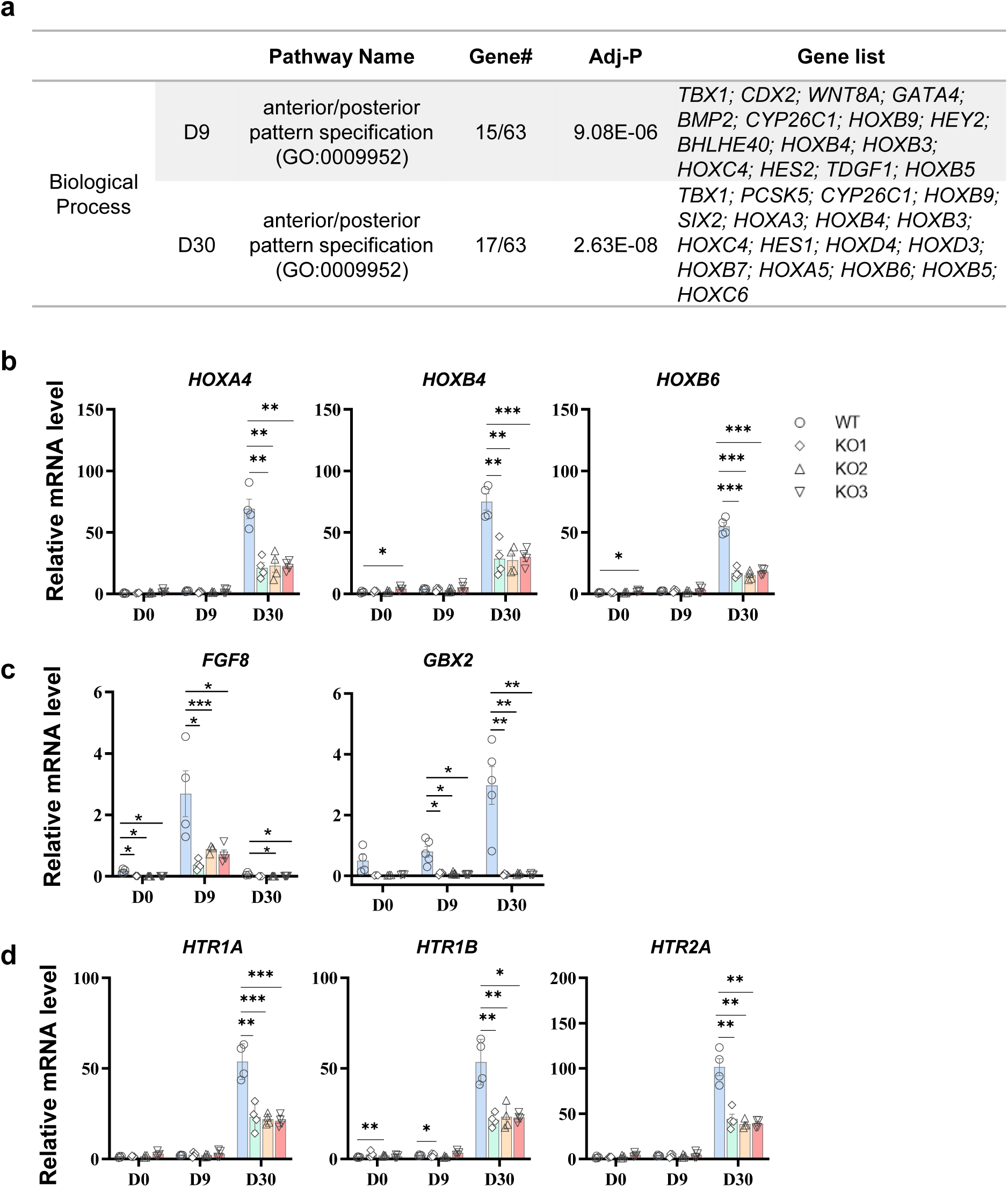
MiR-1202 KO downregulates genes for hindbrain development. a) Pathways enriched for the downregulated DEGs in the miR-1202 KO organoids. Lists of genes in the anterior/posterior pattern specification at D9 and D30 were shown respectively. Gene#, the number of annotated genes in the input list / number of annotated genes in the reference list. *adjP*, *p*-value adjusted by multiple test adjustment. b) Q-PCR analysis of the hindbrain homeobox genes *HOXA4*, *HOXB4* and *HOXB6* were shown. c) Q-PCR analysis of hindbrain patterning factors *FGF8* and *GBX2* were shown. d) Q-PCR analysis of the serotonergic receptor genes HTR1A, HTR1B, HIT2A were shown. Error bars represent mean ± SEM (b, c and d).

### Identification of miR-1202 binding targets

To identify mRNAs that interact with miR-1202, we performed RNA pulldown-seq as described previously (Tan et al., 2014). Biotin-labeled miR-1202 or a control miR (cel-miR-67) were transfected into miR-1202 KO cells at D9 (**Figure 5a**). The levels of miR-1202 were significantly elevated in cells transfected with miR-1202 compared to those transfected with the control miRNA (*p* < 0.0001) (**Figure 5b**). The RNAs captured by biotinylated miR-1202, but not by cel-miR-67, were identified. A total of 697 genes (adjusted *p*-value < 0.05) were recognized as candidate targets of miR-1202-interacting mRNAs from two independent experiments (**Supplementary Table 1**). To narrow down these targets, we compared the candidate genes with the differentially expressed genes (DEGs) that were upregulated in the KO cells (fold change > 2, adjusted *p*-value < 0.05, **Supplementary Table 3**) from the RNA-seq. As shown in the Venn diagram, only 13 genes overlapped by this analysis (**Figure 5c**). We further intersected these datasets with transcripts containing predicted miR-1202 binding sites (a total of 3,895 transcripts identified by the Targetscan algorithm, **Supplementary Table 2**). Through this integrated analysis, we identified five potential targets: *POU3F2*, *ADCYAP1R1*, *TTYH1*, *DCLK2*, and *APC2* (**Figure 5d**).

**Figure 5.**
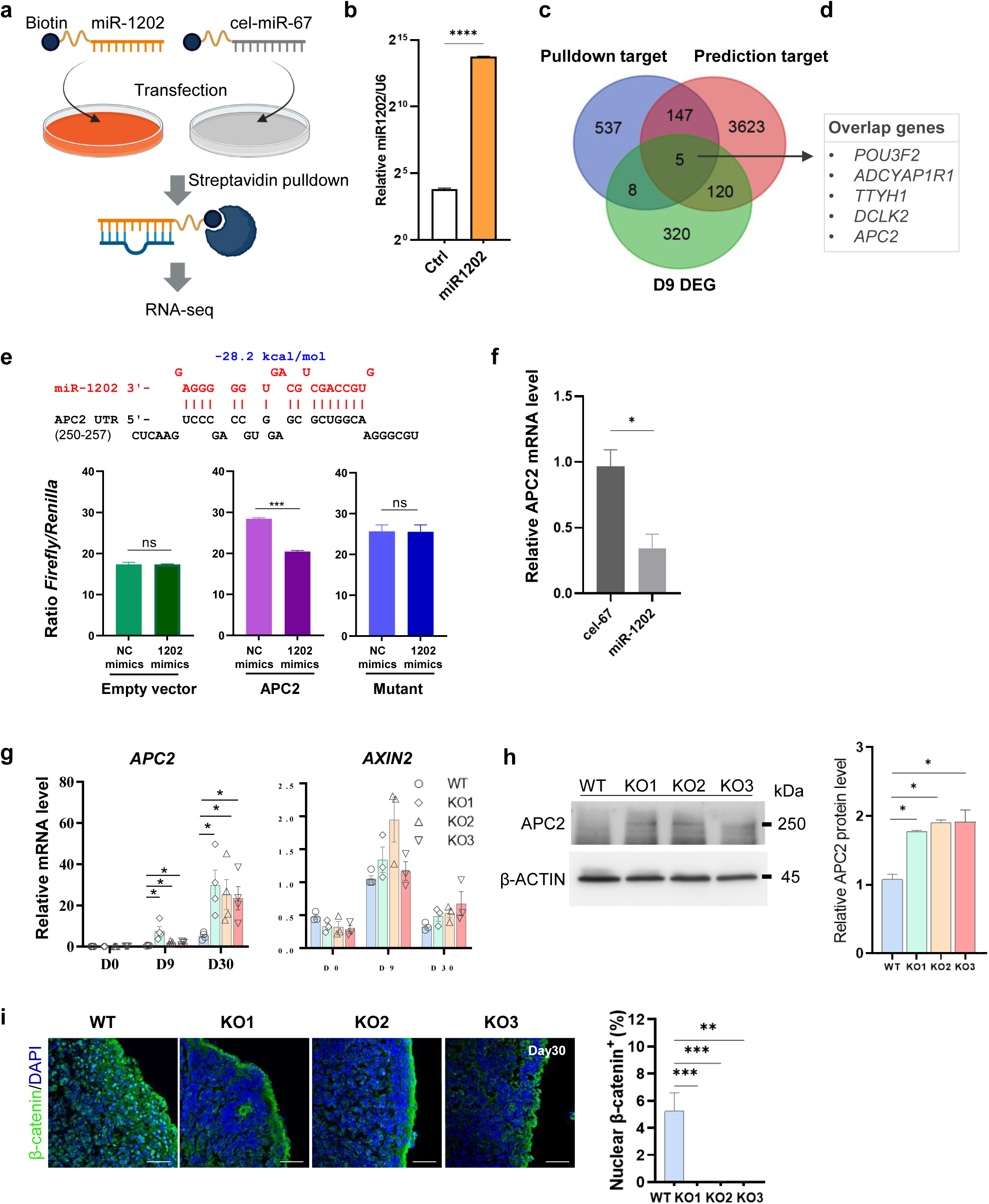
MiR-1202 targets APC2 and modulates WNT signaling during DA neurogenesis. a) Schematic diagram showing RNA pulldown by biotinylated miRNAs followed by RNA-seq for the identification of miR-1202 interacting transcripts. Cel-miR-67 was used as a negative control. b) Q-PCR analysis of the miR-1202 level in the cells transfected with the biotinylated miR-1202 or control miRNA. c) Venn diagram of the overlap genes among pulldown mRNA genes (*adjP* < 0.05), predicted targets (by Targetscan algorithm) and upregulated DEGs at D9 (FC > 2, *adjP* < 0.05). PD: pulldown. d) The list of the overlap genes from c. e) The inhibitive effect of miR-1202 on APC2 3’UTR by luciferase assay. Empty vector (pGL4.13) was used as the negative control. The mutant luciferase plasmid contained the same 3’UTR except that the seed sequence was mutated. The predictive interaction between miR-1202 and APC2 3’-UTR was shown. f) Overexpression of miR-1202 inhibited the APC2 mRNA expression in D9 miR-1202 KO cells. g) Q-PCR analysis of *APC2* and *AXIN2* in WT and miR-1202 KO organoids. h) Western blot analysis of APC2 protein in D44 organoids. Quantification of the protein amount was shown. i) Immunofluorescent staining showing the β-catenin nuclear localization in WT and KO organoids at D30. Quantification of nuclear stains was shown. Scale bars, 500 µm. Error bars represent mean ± SEM (b, e, f, g, h and i).

### MiR-1202 targets APC2 and regulates WNT/**β**-catenin signaling pathway

Among the five potential targets, we were particularly intrigued by *APC2*. APC2 is highly expressed in the brain and acts as a negative regulator of the canonical WNT signaling pathway (Nakagawa et al., 1998). Analysis of the miR-1202 seed binding sequences on the 3’-UTR revealed that APC2 showed high conservation between closely related primates (chimp, gorilla, and orangutan) but not in other animals (**Supplementary Figure 3**). We thus investigated how miR-1202 interacts with *APC2* to regulate DA neurogenesis. A luciferase activity assay was conducted to confirm whether *APC2* is a target of miR-1202. The results demonstrated that miR-1202 mimics could inhibit the luciferase activity; however, there was no inhibition when the seed sequence of the *APC2* 3’UTR was mutated. No significant inhibition was observed in the negative control vector lacking a 3’-UTR (**Figure 5e**). Consistent with the luciferase reporter assay, overexpression of miR-1202 in D9 KO cells suppressed the endogenous *APC2* mRNA levels (**Figure 5f**).

We then examined the gene expressions of *APC2* and *AXIN2* during differentiation. AXIN2 is a downstream transcriptional target of the WNT/β-catenin signaling pathway (AXIN2 is also a cooperative negative regulator with APC2 in this pathway) (Leung et al., 2002). As expected, *APC2* was upregulated in miR-1202 KO cells. *AXIN2* was also upregulated, but showed greater variability among the KO groups (**Figure 5g**). Western blot analysis further confirmed the upregulation of APC2 protein levels in KO organoids compared to WT controls (**Figure 5h**). Since activation of the canonical WNT signaling pathway involves the nuclear localization of β-catenin protein, we assessed β-catenin levels using immunofluorescent staining. As anticipated, nuclear β-catenin signals were significantly reduced in miR-1202 KO organoids (**Figure 5i**). Our results suggest that WNT signaling is downregulated following the knockout of miR-1202 during neural differentiation.

### Knockdown of APC2 in miR-1202 KO organoids attenuated DA neurogenesis

As APC2 negatively regulates WNT/β-catenin signaling by preventing the assembly of the WNT signalosome (Saito-Diaz et al., 2018), we investigated whether the enhanced DA neurogenesis observed in miR-1202 KO cells depended on APC2. To elucidate this mechanism, we generated *APC2* knockdown hPSCs using two different shRNAs (sh1 and sh2) in the KO2 and KO3 clones, respectively. Non-specific control shRNA (shNC) was used for comparison. We then used these four knockdown hPSCs and the two control hPSCs to generate midbrain organoids. The knockdown efficiency of *APC2* in these cells was demonstrated at the mRNA level (D9) and protein level (D44) (**Supplementary Figure 4a,b**). Representative bright-field images of the organoids from each group are shown in **Supplementary Figure 4c**. As expected, the *APC2* knockdowns promoted the growth of organoids from D30 to D44 (**Figure 6a & Supplementary Figure 4d**). Additionally, *APC2* knockdowns rescued the reduced cell proliferation observed in the miR-1202 KO cells, as evidenced by an increased proportion of BrdU^+^ cells and decreased expression of neuron cytoskeleton proteins in APC2-depleted cells (**Figure 6b & Supplementary Figure 4e**).

**Figure 6.**
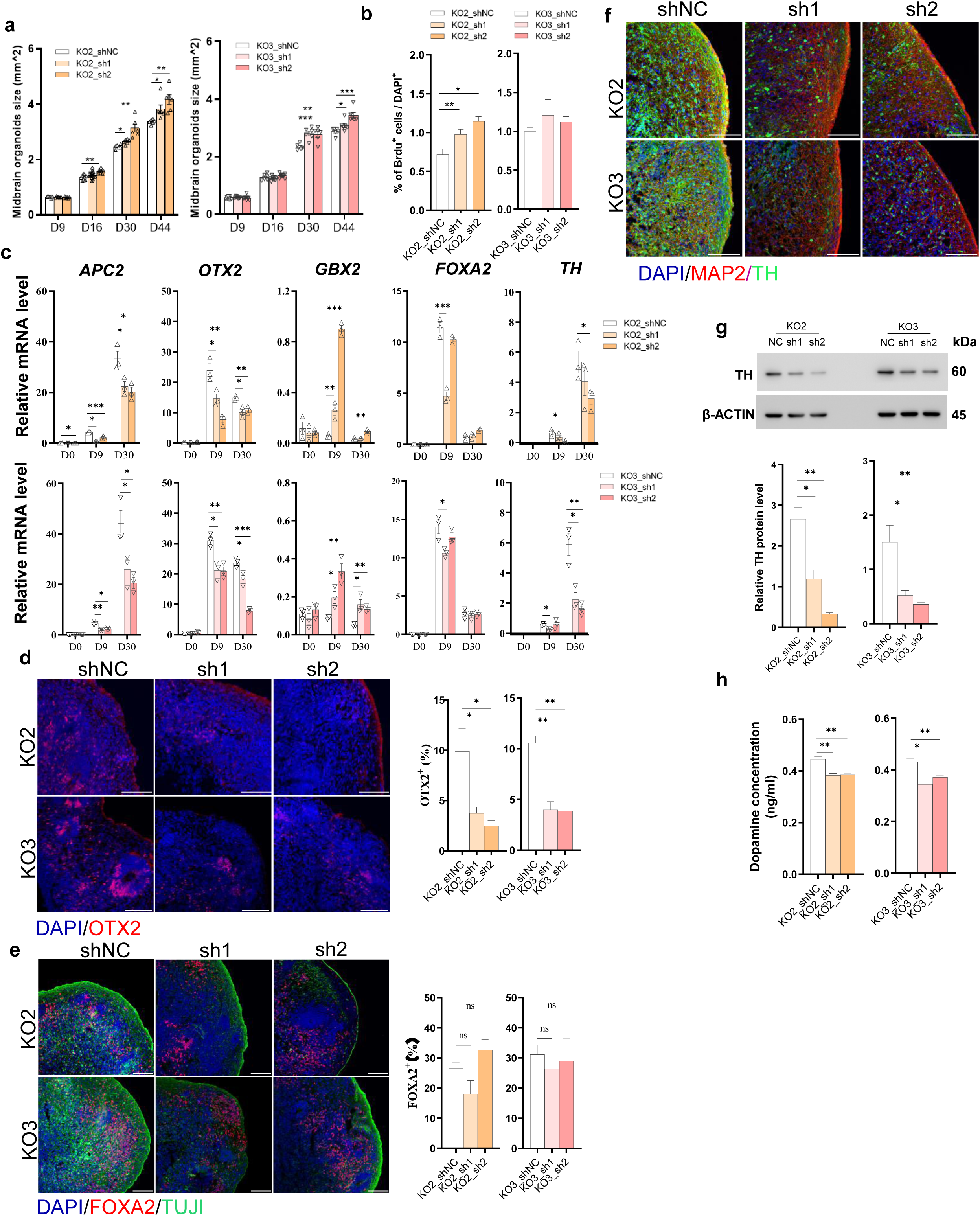
Knockdown of APC2 attenuates DA neurogenesis in miR-1202 KO organoids. a) Stable APC2 knockdown cells were generated by two APC2 shRNAs (sh1 and sh2) in miR-1202 KO2 and KO3 clones. The organoid sizes of each group during differentiation were measured. b) The percentages of BrdU^+^ cells at D30 organoids were measured by flow cytometry. c) Q-PCR analysis of *APC2*, *OTX2*, *GBX2*, *FOXA2* and *TH* expressions from each group of the organoids. d) Immunofluorescent staining showing the positive OTX2 cells at D30 organoids from each group. Quantification was shown. e) Immunofluorescent staining showing the positive FOXA2 and TUJ1 cells at D30 organoids from each group. Quantification was shown. f) Immunofluorescent staining of TH and MAP2 at D65 organoids from each group. g) Western blot analysis of TH protein at D44 organoids from each group. Quantification of protein amount was shown. h) Dopamine levels were measured by LC-MS from each group at D48. Scale bars, 100 µm (d, e and f). Error bars represent mean ± SEM (a, b, c, d, e, g and h).

We analyzed the expression of *APC2* and genes involved in midbrain-hindbrain regionalization (*OTX2* and *GBX2*) as well as DA neurogenesis (*FOXA2* and *TH*) using qPCR. As anticipated, *APC2* mRNA levels decreased with shRNA-mediated knockdown. The midbrain marker *OTX2* showed reduced expression, while the hindbrain marker *GBX2* was increased. *FOXA2* levels were not consistently altered, although *TH* expression consistently decreased in *APC2* knockdown cells (**Figure 6c**). These results were further verified through immunofluorescent staining of OTX2 and FOXA2 (**Figure 6d**). In line with the qPCR results, OTX2 protein levels were significantly downregulated in *APC2* knockdown organoids, whereas FOXA2 protein levels did not show consistent changes (**Figure 6e**). To confirm the decrease in TH at the protein level, we performed immunofluorescent staining and Western blot analysis (**Figure 6f,g**). Both analyses consistently indicated lower TH levels in *APC2* knockdown organoids. Finally, we measured dopamine levels using LC-MS and found decreased dopamine levels in the *APC2* knockdown groups (**Figure 6h**). Together, these results support that miR-1202 negatively regulates APC2 to suppress DA neuron differentiation.

### MiR-1202 and APC2 are responsive to antidepressant treatments

MiR-1202 is known to be a diagnostic biomarker for MDD, with remitters demonstrating increased levels of miR-1202 after eight weeks of antidepressant treatment (Lopez et al., 2014). Common prescribed antidepressants, such as citalopram and imipramine, are selective serotonin reuptake inhibitors that prevent the reuptake of neurotransmitters by presynaptic neurons (Rosenberg et al., 1994). In our study, we investigated whether citalopram and imipramine could stimulate miR-1202 expression and influence DA neurogenesis.

We treated neural progenitor cells with citalopram and imipramine for 24 hours to simulate acute treatment, as well as for 14 days to mimic chronic treatment, using nontoxic concentrations (**Supplementary Figure 5**). We revealed that miR-1202 expression was upregulated in both acute and chronic treatments, with citalopram resulting in a higher level of upregulation compared to imipramine (**Figure 7a, b**). Additionally, we observed a decrease in *APC2* expression following drug treatments (*p* < 0.05 for both citalopram and imipramine) (**Figure 7c, d**). Midbrain markers, such as *OTX2* and *LMX1A*, were also reduced. Conversely, *GBX2* expression increased with the drug treatments. Notably, more significant changes were observed with chronic treatment (**Figure 7c, d**). These results suggest that antidepressants may inhibit DA neurogenesis while promoting serotonergic differentiation in response to treatment. Lastly, we explored whether APC2 is associated with MDD by comparing its expression in MDD patients (n = 9) and non-psychiatric healthy controls (n = 29) using postmortem brain samples from a public dataset (GSE101521) (Pantazatos et al., 2017). We found a significant upregulation of *APC2* mRNA in MDD patients compared to healthy controls (**Figure 7e**, *p* < 0.01). As both miR-1202 and *APC2* are enriched in the brain and linked to MDD, our study highlights a previously unknown regulatory axis (miR-1202/APC2/WNT) that may play a crucial role in fine-tuning neuronal fate in the brain (**Figure 7f**).

**Figure 7.**
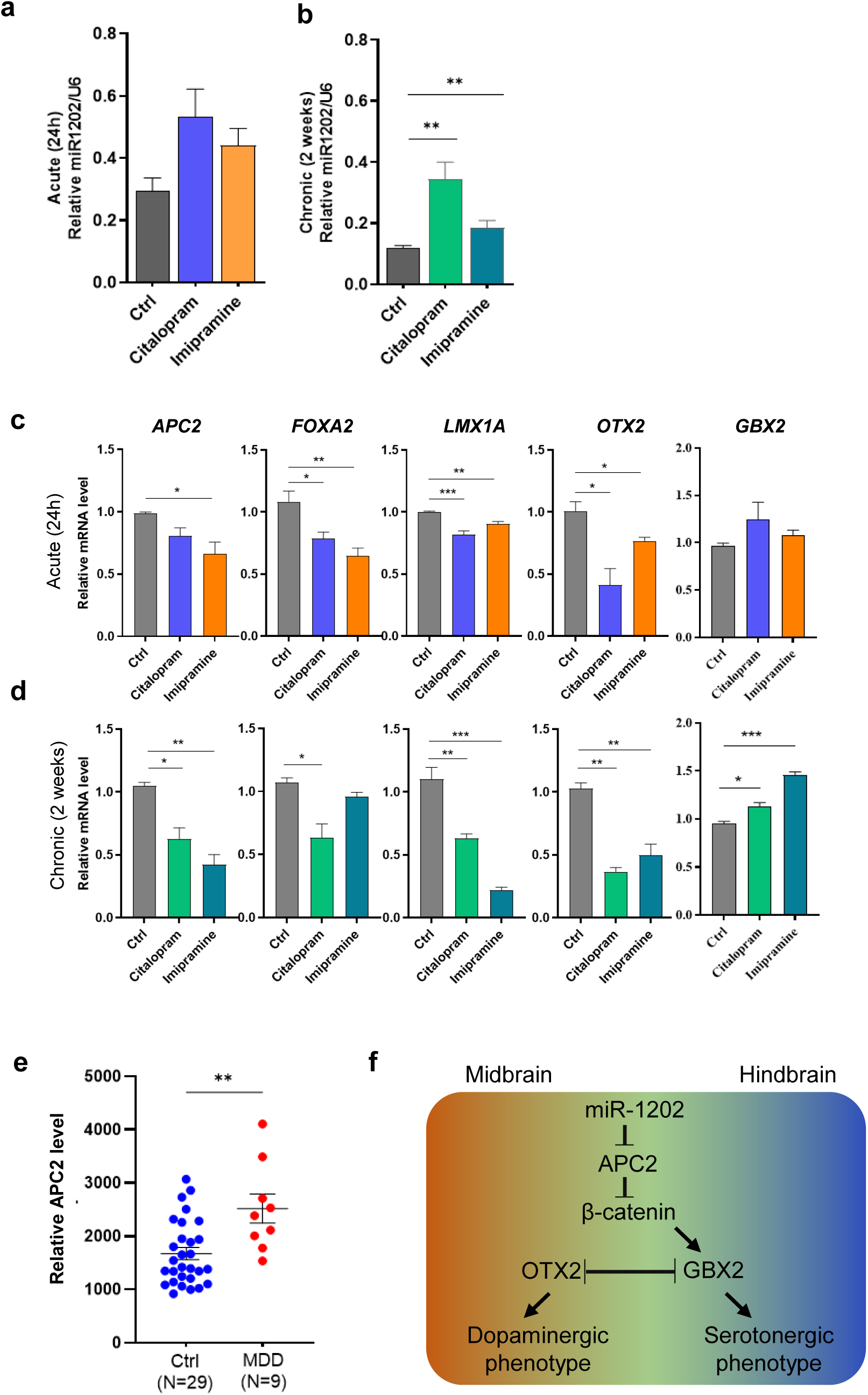
MiR-1202 and APC2 respond to antidepressant treatments and the effect of the drugs on brain organoid differentiation. a, b) MiR-1202 expressions in cells treated with antidepressant drugs (50 µM for citalopram and 12.5 µM for imipramine) for 24 hours (acute treatment) and 2 weeks (chronic treatment). c, d) Q-PCR analysis of *APC2*, *FOXA2*, *LMX1A*, *OTX2* and *GBX2* expressions after acute and chronic treatments with the antidepressant drugs. e) *APC2* expressions in the postmortem brain in MDD patients (N = 9) and non-psychiatric healthy controls (N = 29). Data were retrieved from public dataset (GSE101521). f) Schematic diagram summarizing the regulatory relationship between miR-1202 and APC2 and the downstream β-cateinin signaling that fine-tunes the neuronal differentiation in the midbrain-hindbrain boundary. Error bars represent mean ± SEM (a, b, c and d).

## DISCUSSION

MiR-1202 is unique to primates, and suitable animal models for studying primate-specific genes or miRNAs are not readily available. To mimic brain development, we employed a human brain organoids model (Jo et al., 2016). Deletion of miR-1202 resulted in reduced organoid growth, accompanied by decreased cell proliferation. Notably, microdeletions at 6q25.2-25.3, where miR-1202 is mapped to, have been observed in patients with microcephaly and developmental delays (Lukusa et al., 2001; Nagamani et al., 2009). RNA-seq revealed that DA neurogenesis was upregulated in the miR-1202 KO organoids. Encouraged by these findings, we further explored the molecular basis underlying miR-1202-mediated gene regulation. Using the RNA pulldown-seq approach, we identified several candidate genes that negatively correlated with miR-1202 and contained miRNA binding sites, focusing particularly on *APC2*, which is also enriched in the brain. A luciferase reporter assay confirmed the inhibitory effects of miR-1202 on the 3’-UTR of *APC2*. APC2 functions as a negative regulator of the canonical WNT signaling pathway (Nakagawa et al., 1998), and its upregulation decreases WNT signaling. APC2 has been known to be regulated by other miRNAs such as miR-100 and miR-125b (Lu et al., 2017). Furthermore, we demonstrated that nuclear β-catenin signaling was significantly decreased in miR-1202 KO organoids. WNT signaling plays temporally distinct roles in early brain development (Bem et al., 2019; Mulligan and Cheyette, 2012). It has been reported that strong WNT/β-catenin signaling inhibits the differentiation of mesodiencephalic progenitors into Pitx3^+^ and Th^+^ mdDA neurons by repressing *Pitx3* in mice (Nouri et al., 2020). Consistent with these reports, reduced β-catenin signaling in miR-1202 KO cells promoted differentiation into DA neurons.

GBX2, a direct target activated by WNT/β-catenin signaling (Li et al., 2009), interacts with OTX2 to mutually repress the expressions of the counterparts at the midbrain-hindbrain boundary (Harada et al., 2016). OTX2 is a transcription factor critical for neurogenesis and the specification of ventral midbrain DA neurons (Omodei et al., 2008; Puelles et al., 2004), whereas GBX2 opposes OTX2 and promotes hindbrain development (Millet et al., 1999). Our results indicated that lower β-catenin signaling in miR-1202 KO cells downregulated *GBX2* expression, subsequently increasing *OTX2* expression and enhancing DA neurogenesis.

The primary population of DA neurons in the mammalian brain resides in the ventral midbrain. An earlier study examined the effects of WNT upregulation on midbrain DA neuron development, demonstrating that activated WNT signaling not only downregulates *SHH* but also *FOXA2* (Nouri et al., 2015). In our study, we also noted the upregulation of *SHH* and *FOXA2* in the miR-1202 KO organoids during differentiation. These findings aligned with the weakened WNT/β-catenin signaling, resulting in enhanced DA neurogenesis.

Depression and other psychiatric disorders often result from impaired communication between presynaptic and postsynaptic neurons. Tricyclic antidepressants work by preventing the reabsorption of neurotransmitters, thereby increasing their availability in the synapse (Plenge et al., 2020; Sindrup et al., 2005). In patients with MDD, the microRNA miR-1202 is dysregulated. This led us to investigate whether miR-1202 and its downstream target, APC2, respond to antidepressant treatments and how these drugs affect dopaminergic phenotype. Our findings showed that chronic treatment resulted in the upregulation of miR-1202, while also leading to decreased expression of *APC2* and other markers associated with dopaminergic neurogenesis such as *FOXA2*, *LMX1A*, and *OTX2*. However, the physiological significance of such changes in response to antidepressant treatment cannot be evaluated by the current cell model, which is one of the limitations. How miR-1202 mediates antidepressant treatment may require a further study in primate model.

In summary, we employed an organoid model that permitted us to study the function of miR-1202 to mimic the developing brain. Loss of miR-1202 repressed organoid growth and promoted DA neuron differentiation. MiR-1202 appeared to suppress *APC2*, and thus, loss of miR-1202 inhibited β-catenin signaling to modulate the DA neurogenesis. This effect was abrogated by knocking down *APC2*. We also demonstrated that both miR-1202 and *APC2* responded to antidepressants, implying a potential role of this regulatory axis in psychiatric disorders.

## MATERIALS AND METHODS

### Cell culture

The hPSC line WA01 (H1) was obtained from WiCell. H1 cells were maintained in mTeSR^TM^-Plus medium (STEMCELL Technologies). HEK293T cell line was obtained from ATCC. HEK293T cells were cultured in DMEM medium (Thermofisher) supplemented with 1% GlutaMAX (Thermofisher), 10% fetal bovine serum (FBS) (Thermofisher) and 1% penicillin/streptomycin (PS) (Thermofisher).

### Generation of miR-1202 knockout cell

To generate miR-1202 KO cells, H1 cells were transfected with a pair of CRISPR-Cas9 plasmids (PX458, Addgene# 48138) that targeted the upstream and downstream sites of the mature miR-1202 genomic sequence. Cells after nucleofection were GFP-sorted, and allowed to expand to mixed colonies (KO1) or single cell-derived colonies (KO2 and KO3). Control (WT) cells were generated by the same procedure but with a non-targeting gRNA sequence. Cells were characterized by expressions of pluripotency markers and Sanger sequencing of the targeted locus.

### Generation of APC2 knockdown cell

All shRNAs used in this study were cloned into pDECKO_mCherry lentivector (Addgene#78534) and packaged into pseudo-lentiviral particles. The H1 cells were maintained at ∼50% confluency in a single 6-well plate and infected with 20 μl lentiviral particles (MOI = 1) in 2 mL cell culture medium supplemented with 6 µg/ml polybrene (Sigma-Aldrich). Transduced cells were sorted by FACS and expanded. The shRNAs for APC2 are listed below: Sh1: 5’-GTACTGCCCACGCGAACATAT; and Sh2: 5’-CCAGATGAGGTGAAAGCTTAT.

### Generation of midbrain organoids

Midbrain organoids were generated as previously described (Hou et al., 2022; Jo et al., 2016). Briefly, hPSCs were seeded in the ultra low–attachment U-bottom 96-well plates (1 x 10^4^ cells per well) to generate uniform embryonic bodies (EBs) with EB medium (D0). The EB medium was composed of DMEM/F12 (Thermofisher) supplemented with 20% Knockout Serum Replacement (Thermofisher), 1% PS, 1% GlutaMAX (Thermofisher), 1% NEAA (Thermofisher), 55 μM β-mercaptoethanol (Thermofisher), 1 μg/ml of heparin (STEMCELL Technologies), 3% FBS (Thermofisher), 4 ng/ml of bFGF (Peprotech), and 50 μM Y27632 (Peprotech). After 24 h (D1), the medium was switched to brain organoid generation medium (BGM) supplemented 2 μM dorsomorphin (Sigma-Aldrich), 2 μM A83-01 (Peprotech), 3 μM of CHIR99021 (Peprotech) and 1 μM IWP2 (Peprotech) and changed every other day. The BGM contained 1:1 mix of DMEM/F12 and Neurobasal Medium (Thermofisher), 1x N2 supplement (Thermofisher), 1x B27 without vitamin A (Thermofisher), 1% PS, 1% GlutaMAX, 1% NEAA, 55 μM β-mercaptoethanol and 1 μg/ml heparin. On D4, 100 ng/ml FGF8 (Peprotech) and 2 μM SAG (Peprotech) were added to induce specification into the mesencephalic floor plate. After 3 days, organoids were embedded into growth factor-reduced matrigel (Corning) droplets. Matrigel-embedded MBOs were transferred onto 6-cm petri dishes containing BGM supplemented with 100 ng/ml of FGF8, 2 μM SAG, 200 ng/ml laminin (Sigma-Aldrich), and 2.5 μg/ml insulin (Thermofisher). After 2 days, MBOs were transferred into ultra low–attachment 6-well plates (Corning) containing brain organoid maturation medium (BMM), consisting of 1:1 mix of DMEM/F12 and Neurobasal Medium supplemented with 1x N2 supplement, 1x B27 without vitamin A, 1% PS, 1% GlutaMAX, 1% NEAA, 55 μM β-mercaptoethanol, 1 μg/ml heparin, 10 ng/ml BDNF (Peprotech), 10 ng/ml GDNF (Peprotech), 200 μM ascorbic acid (Sigma-Aldrich) and 125 μM db-cAMP (Peprotech). From this maturation step, the organoids were cultured on an orbital shaker (Thermofisher). BMM was replaced every other day.

### Gene expression analysis

Total RNA and miRNA were extracted using *mir*Vana™ miRNA Isolation Kit (Thermofisher) according to the instruction. Total mRNA was reversed transcribed using PrimeScript RT-PCR Kit (Takara). Quantitative RT-PCR (qPCR) was performed with the SYBR green PCR Master Mix (Thermofisher) using the ABI Quantstudio 7 Pro Real-time PCR system (Thermofisher). ΔCt values were calculated by subtracting the Gapdh Ct value from that of each target gene. Relative expression levels were calculated by using 2^−ΔΔCt^ methods. Primer sequences used for qPCR are listed in Supplementary Table 4. miRNA was reverse transcribed using TaqMan RT-PCR microRNA assays (Thermofisher) according to the manufacturer’s instructions. For miRNA quantification, miRNA was normalized to U6.

### BrdU-based cell proliferation assay

At D30, organoids were incubated with culture medium supplemented with10 µM BrdU for 2[h. The organoids were harvested and washed with PBS three times, and then dissociated into single cells. The cell proliferation rates were detected by flow cytometry according to the instruction.

### LC-MS

Conditioned culture medium was collected after an overnight exposure at the D48 differentiated organoids. Acetonitrile was used to precipitate proteins out of the collected medium. After that, the supernatant was filtered and subsequently analyzed using Agilent 6460 Triple Quadrupole LC-MS system. The dopamine standard was purchased from Sigma-Aldrich. The culture medium conditioned in empty well was used as negative control.

### RNA sequencing

The extracted total RNA for sequencing was performed by Beijing Novogene Technology Co. Ltd. DEGs were subjected to GO and Wikipathway enrichment analysis using EnrichR online software and EdgeR algorithm.

### Western blot

Total proteins were isolated from cells or organoids by RIPA buffer, and protein concentrations were measured by DC protein assay method of Bradford (Bio-Rad). Five microgram of denatured protein from each sample were separated on 10% SDS-PAGE and then transferred onto PVDF membranes. After blocking with 5% bovine serum albumin (BSA), blots were incubated with primary antibodies overnight at 4°C and secondary antibodies for 1 h at room temperature respectively. Membranes exposed to ECL western blotting substrate (Thermofisher). Band intensities were determined by Image J (NIH). Primary antibodies used for Western were listed as follows: TH (Sigma-aldrich), APC2 (Sigma-aldrich) and β-actin (CST).

### Immunofluorescence

The organoids were fixed in 4% paraformaldehyde (PFA) (Sigma-aldrich), washed in PBS, incubated in a 30% sucrose solution in PBS at 4°C overnight, and subsequently embedded in O.C.T compound (SAKURA) for cryosectioning. The organoids were cryosectioned at a thickness of 10μm using Cyrost (Leica CM1950). The slices were blocked with 5% BSA with 0.1% Triton X-100 in PBS for 1 h at room temperature, incubated with primary antibodies overnight at 4°C, and secondary antibodies for 1 h at room temperature. All slices were counterstained with DAPI (Sigma-Aldrich) and mounted with Prolong gold antifade mounting media (Thermofisher). Image was taken on a Leica SP8 confocal microscope (Leica). Primary antibodies used for immunofluorescence were listed as follows: SOX2 (CST), NESTIN (CST), FOXA2 (Santa Cruz), TUJ1 (Abcam), OTX2 (Merck millipore), KI67 (CST), MAP2 (Abcam), TH (Sigma) and β-CATENIN (Thermofisher).

### Biotin RNA pulldown and RNA-seq

Biotin pulldown assay were performed as described (Tan and Lieberman, 2016). In brief, 1 × 10^6^ cells dissociated from D9 organoids were seeded on Geltrex (Thermofisher) coated 6-well plate and cultured overnight at 37°C in a 5% CO2 incubator. The cultured cells were transfected with 100 picomoles of biotinylated miR-1202 and cel-miR-67 miRNA mimics (Genepharm) separately using Lonza Nucleofector device according to the instruction. Biotin was attached to the 3’-end of the active strand. The cell pellets were collected 24 h post transfection, washed twice with cold PBS, and incubated with lysis buffer [20 mM Tris (pH 7), 25mM EDTA (pH 8), 100 mM NaCl, 5 mM MgCl_2_ (all Thermofisher), 2.5 mg/ml Ficoll PM400, 7.5 mg/ml Ficoll PM70, 0.25 mg/ml dextran sulfate 670k, 0.5% IGEPAL (Sigma-Aldrich), 50 U RNase OUT and 50U SUPERase In (Thermofisher), and complete protease inhibitor cocktail (Roche)] on ice for 20 min. The cytoplasmic lysate, isolated by centrifugation at 5,000 g for 5 min, was added to 1 mg/ml yeast tRNA (Thermofisher) and 1 mg/ml BSA-blocked streptavidin (SA)-coated magnetic beads (Thermofisher) and rotated for 4 h at 4°C. The beads were washed five times with 1 ml lysis buffer and bead-bound RNA was extracted by *mir*Vana™ miRNA Isolation Kit. Libraries were constructed by using NEBNext Single Cell/Low Input RNA Library Prep Kit for Illumina (NEB E6420S) according to the manufacturer’s instructions.

### Dual luciferase reporter assay

HEK293T cells were grown in 96-well plates and transfected with 10[nM *mir*Vana™ miR-1202 mimics (Thermofisher, #4464084), 100 ng of pGL4.13 vector (Promega) tagged with the 3′-UTR that includes miR-1202-binding sites and pRL Renilla Luciferase control reporter vector (Promega). *mir*Vana™ miRNA Mimic, Negative Control #1 (Thermofisher, #4464058) was used as the negative control. The Firefly and Renilla luciferase activities in the cell lysates were assayed with a Dual-Luciferase Reporter Assay System (Promega) at 48 h post-transfection according to the instruction.

### Antidepressant treatment

The D9 midbrain organoid cells were used for screening for cytotoxic effects by cell-counting kit-8 (DOJINDO) according to the manufacturer’s protocol. The antidepressants were applied at nontoxic concentrations. Cells were grown in the continuous presence of either 50 μM citalopram hydrobromide (MCE), 12.5 μM imipramine hydrochloride (MCE) or vehicle control for 24 h (acute) or 14 d (chronic).

### Statistical analysis

For statistical analysis, the unpaired *t*-test and *ANOVA* were used for calculating *p* values. The *p* values were presented as * *p* < 0.05; ** *p* < 0.01; and *** *p* < 0.001.

## Supplemental Information

### Supplementary Figure Legends

**Supplementary Figure 1.** Generation of miR-1202 KO cells in hPSC. a) Sanger sequencing confirmed the deletion of the mature miR-1202 sequence and the upstream promoter. b) Genomic PCR products showing shortened DNA fragments after deletion.

**Supplementary Figure 2.** Q-PCR analysis of additional DA neuronal markers (ASCL1, NR4A2, NEUROG2), and pan-neuronal marker (TUBB3) in WT and miR-1202 KO organoids during differentiation. Error bars represent mean ± SEM.

**Supplementary Figure 3.** Alignments of the miR-1202 seed sequence (UGCCAGC) with the 3’ UTR of the identified target genes: (a) APC2, (b) POU3F2, (c) ADCYAP1R1, (d) TTHY1, (e) DCLK2. Sequence conservation among different animal species were shown (solid box). (f) Gene expression of APC2 in different human tissues. Data was retrieved from GTEx Portal (https://www.gtexportal.org/).

**Supplementary Figure 4.** a) Stable APC2 knockdown PSC lines were generated by two APC2 shRNA (sh1 and sh2) and negative control shRNA (shNC) in miR-1202 KO2 and KO3 cells, respectively. Q-PCR analysis showed the knockdown efficiencies of the shRNAs at D9 organoids. b) Western blot analysis showed the knockdown efficiencies of APC2 at D44. c) Representative bright-field images showing the growth of the developing organoids from the six stable lines. d) Quantification of the organoid weights during differentiation. e) The relative mRNA levels of pan-neuron makers (MAP2 and TUBB3) were measured by q-PCR. Error bars represent mean ± SEM.

**Supplementary Figure 5.** Drug toxicity of citalopram and imipramine were tested at different concentrations (Citalopram: 0 µM, 50 µM, 100 µM, and 200 µM; Imipramine: 0 µM, 6.25 µM, 12.5 µM, and 25 µM) using the CCK8 cytotoxicity assay. Error bars represent mean ± SD.

### Supplementary Tables

**Supplementary Table 1.** The list of potential targets identified as miR-1202 interacting mRNAs by pulldown RNA sequencing (*adjP* < 0.05).

**Supplementary Table 2.** The transcripts with miR-1202 interacting sites predicted by Targetscan algorithm.

**Supplementary Table 3.** The upregulated DGEs list among WT and three KOs lines from RNA sequencing at D9 (fold change > 2, *adjP* < 0.05)

**Supplementary Table 4.** Primers used in this study.

## Acknowledgements

The work described in this paper was supported by a grant from the Research Grants Council of the Hong Kong Special Administrative Region, China (Project No. 14169717) and a grant from Health@InnoHK Program launched by Innovation Technology Commission of the Hong Kong Special Administrative Region, China.

## Author contributions

XL performed the majority of the experiments, analyzed the data and drafted the manuscript. LL, SH and DC performed some of the experiments. SWC performed the bioinformatics analysis. WYC and HHC supervised the study. HHC designed and led the project and drafted the manuscript.

## Data availability

The RNA-seq data are deposited in NCBI with the accession code PRJNA894813.

