## Supplementary Figures for "Primate-specific microRNA-1202 regulates dopaminergic neurogenesis by targeting APC2 and modulates WNT/β-catenin signaling pathway in midbrain organoid"

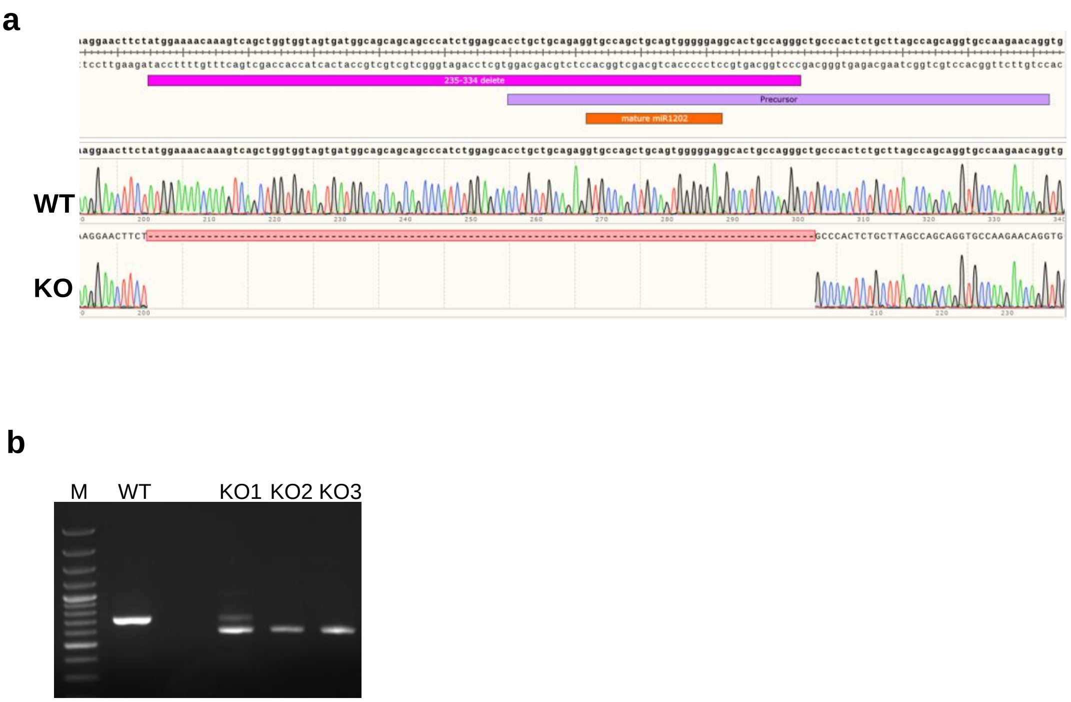


**Supplementary Figure 1. Generation of miR-1202 KO cells in hPSC.** a) Sanger sequencing confirmed the deletion of the mature miR-1202 sequence and the upstream promoter. b) Genomic PCR products showing shortened DNA fragments after deletion. KO1: heterogeneous KO clones; KO2 and KO3: single cell-derived clones.

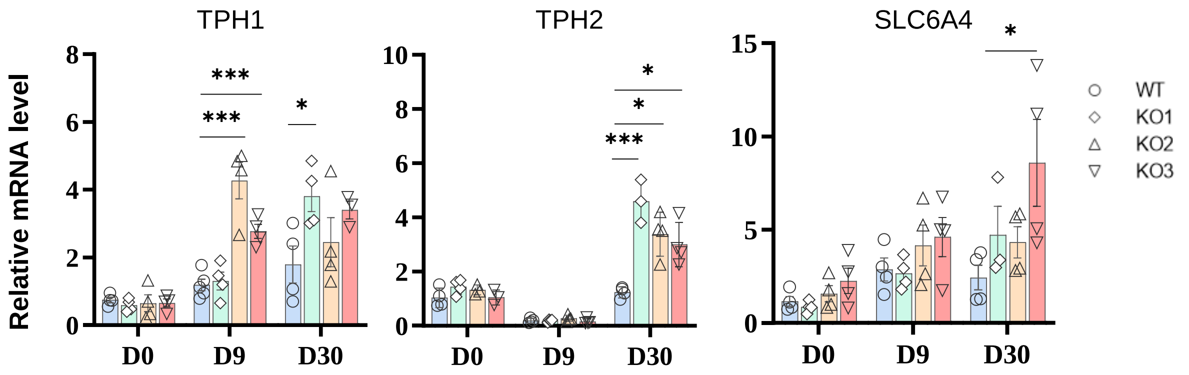


**Supplementary Figure 2.** Q-PCR analysis of serotine synthesis enzymes TPH1 and TPH2 and serotine neurotransmitter transporter SLC6A4 in WT and miR-1202 KO organoids during differentiation. Error bars represent mean ± SEM.

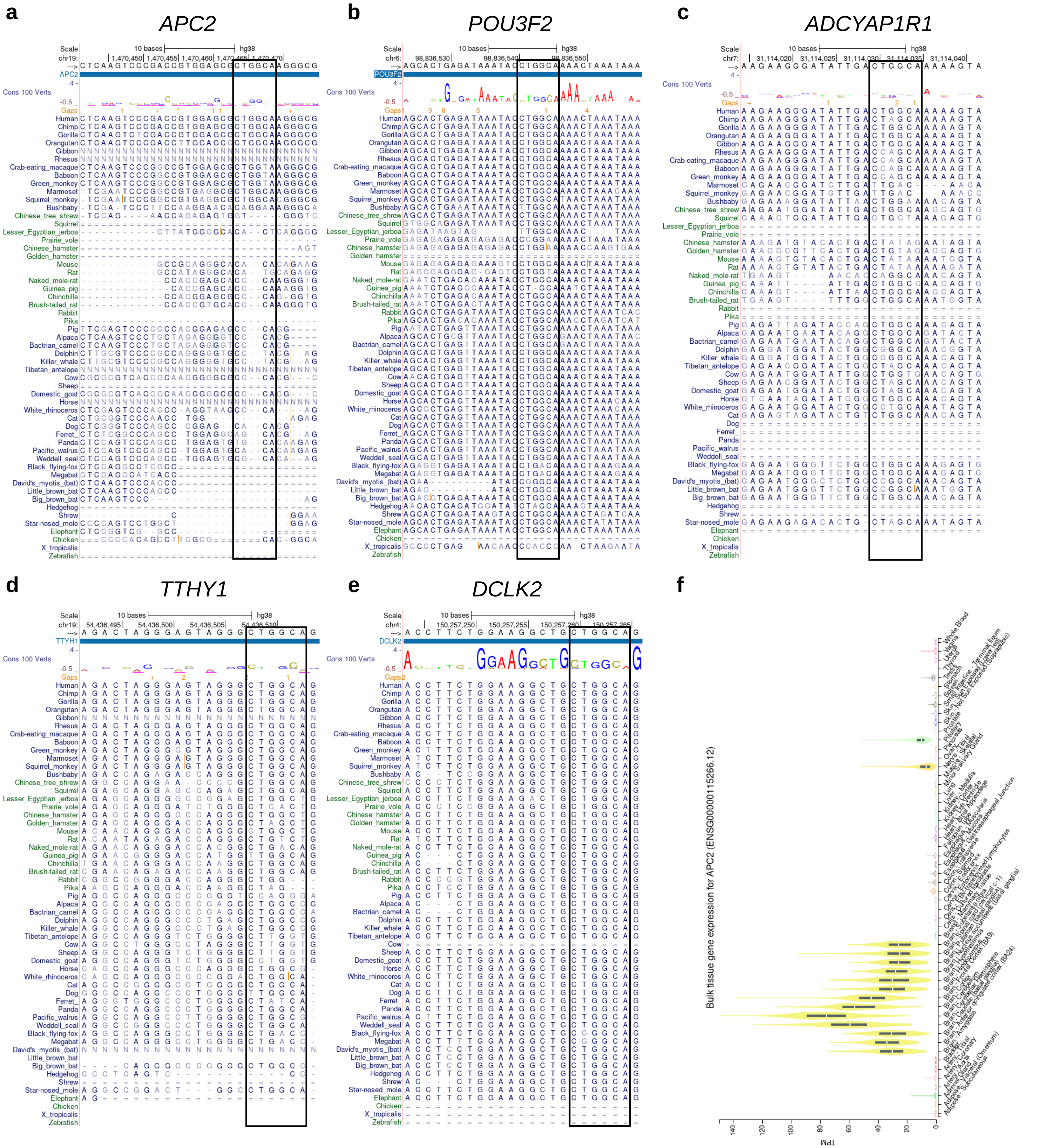


**Supplementary Figure 3.** Alignments of the miR-1202 seed sequence (UGCCAGC) with the 3' UTR of the identified target genes: (a) *APC2*, (b) *POU3F2*, (c) *ADCYAP1R1*, (d) *TTHY1*, (e) *DCLK2*. Sequence conservation among different animal species were shown (solid box). (f) Gene expression of APC2 in different human tissues. Data was retrieved from GTEx Portal (https://www.gtexportal.org/).

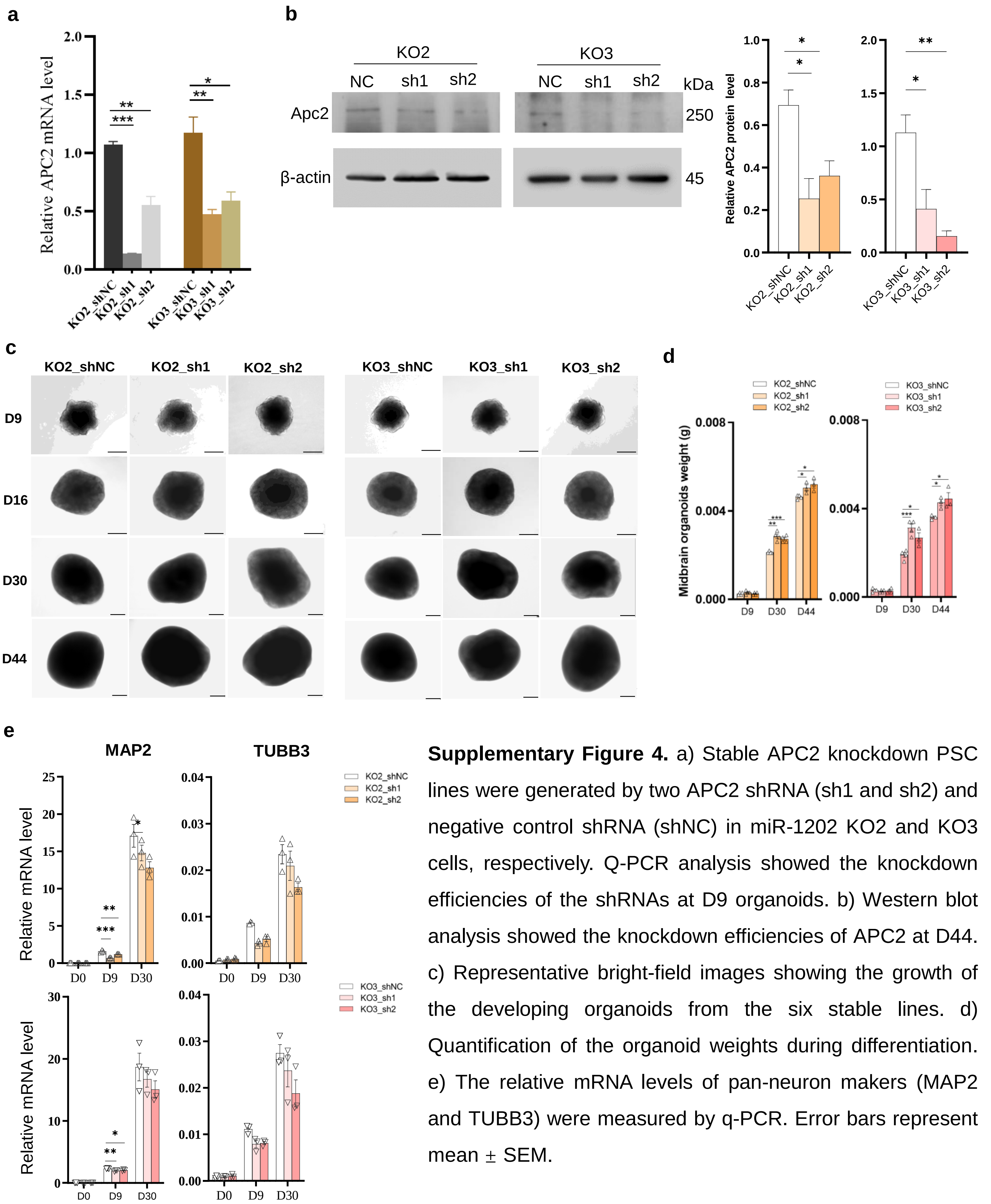


**Supplementary Figure 4.** a) Stable APC2 knockdown PSC lines were generated by two APC2 shRNA (sh1 and sh2) and negative control shRNA (shNC) in miR-1202 KO2 and KO3 cells, respectively. Q-PCR analysis showed the knockdown efficiencies of the shRNAs at D9 organoids. b) Western blot analysis showed the knockdown efficiencies of APC2 at D44. c) Representative bright-field images showing the growth of the developing organoids from the six stable lines. d) Quantification of the organoid weights during differentiation. e) The relative mRNA levels of pan-neuron makers (MAP2 and TUBB3) were measured by q-PCR. Error bars represent mean ± SEM.

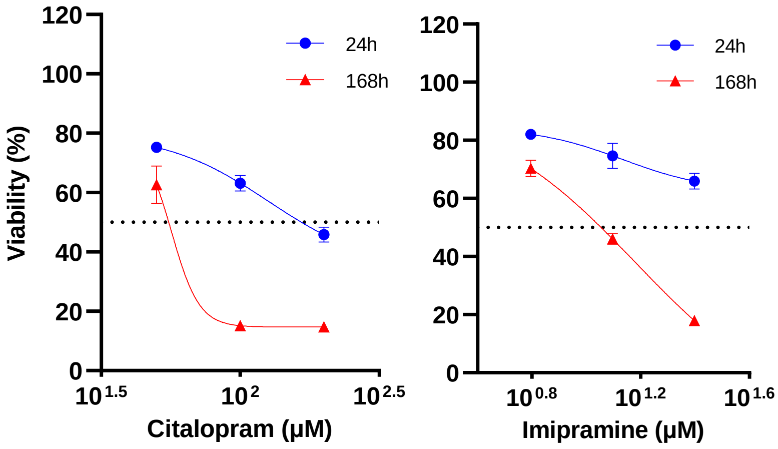


**Supplementary Figure 5.** Drug toxicity of citalopram and imipramine were tested at different concentrations (Citalopram: 0 μM, 50 μM, 100 μM, and 200 μM; Imipramine: 0 μM, 6.25 μM, 12.5 μM, and 25 μM) using the CCK8 cytotoxicity assay. Error bars represent mean ± SD.
